# Improving Protein Interaction Prediction in GenPPi: A Novel Interaction Sampling Approach Preserving Network Topology

**DOI:** 10.1101/2024.08.16.608332

**Authors:** Alisson Silva, Carlos Marquez, Iury Godoy, Lucas Silva, Matheus Prado, Murilo Beppler, Natanael Avila, Anderson R. Santos

## Abstract

**Background:** Computational prediction of protein-protein interactions (PPIs) is crucial for understanding cell biology and drug development, offering an alternative to costly experimental methods. The original GenPPi software advanced *ab initio* PPI network prediction from bacterial genomes but was limited by its reliance on high sequence similarity. This work introduces GenPPi 1.5 to enhance these predictive capabilities.

**Results:** GenPPi 1.5 incorporates a Random Forest (RF) algorithm, trained on 60 biophysical features from amino acid propensity indices, to classify protein similarity even in low sequence identity scenarios (targeting ***>***65% identity). To manage computational complexity from the increased interactions generated by the RF model, especially in extensive conserved phylogenetic profiles, we developed and integrated the Reduced Interaction Sampling (RIS) algorithm. RIS stochastically samples interactions within these profiles, optimizing performance for complete genome analysis. Extensive simulations across various configurations validated the methodology. RF integration significantly broadened GenPPi’s predictive power; application to *Buchnera aphidicola* showed up to 62% overlap with STRING database interactions. Analysis of RIS demonstrated that while introducing some randomness, critical node identification remains robust, particularly for Top N values ***≥*** 100, indicating minimal compromise to network integrity.

**Conclusion:** The combination of Machine Learning (RF) and the RIS algorithm in GenPPi 1.5 represents a significant advancement. It overcomes the high-similarity dependency of the previous version while efficiently handling complex genomes. GenPPi 1.5 provides a robust and scalable alignment-free PPI prediction solution, enabling users to train custom models tailored to specific genomic contexts. GenPPi is freely available on our website https://genppi.facom.ufu.br/, its source code is hosted on GitHub https://github.com/santosardr/genppi, and it can be easily installed via the Python Package Index using the command pip install genppipy.

## 1 Introduction

Protein-protein interactions (PPIs) orchestrate fundamental cellular processes, making their identification crucial for understanding biological systems and developing therapeutic strategies [1]. While experimental methods provide valuable data, they are often costly and time-consuming [2], motivating the development of computational prediction approaches [1]. However, a significant challenge arises when analyzing the vast number of newly sequenced or poorly annotated genomes, mainly from bacteria and archaea. Many computational methods rely heavily on existing experimental data, structural information, or extensive functional annotations (e.g., [3, 4]), limiting their applicability precisely where discovery potential is high in less-characterized organisms [5]. While methods leveraging protein sequence homology (interologs) or 3D structural information [3, 6] can be powerful, they are often limited by the availability of well-annotated homologs with known interactions or experimentally determined/accurately predicted structures, respectively. This is a particular bottleneck for novel proteins from newly sequenced genomes.

Addressing this gap requires robust *ab initio* methods capable of predicting PPI networks directly from genomic sequences with minimal reliance on external databases beyond sequence repositories. The GenPPi software was initially developed with this goal, utilizing genomic context features like conserved gene neighborhoods, phylogenetic profiles, and gene fusion events. We developed the program to analyze genomes represented in multiple files of protein sequences. [5]. While representing an advance, the original GenPPi (v1.0) faced a critical limitation: its protein similarity assessment required a high sequence identity (*>* 90%), significantly restricting its sensitivity and potentially missing biologically relevant interactions between more divergent proteins. Furthermore, simply lowering this threshold introduces a secondary challenge: the combinatorial explosion of potential interactions, which can overwhelm downstream analyses.

This work presents GenPPi 1.5, designed to overcome this dual challenge by integrating a sensitive Random Forest (RF) model for similarity assessment with the novel Reduced Interaction Sampling (RIS) algorithm to manage computational complexity. Computational approaches to PPI prediction encompass a diverse range of methodologies [1]. Structure-based methods leverage 3D protein coordinates for docking or interface analysis but require known or accurately predicted structures, which are often unavailable [3, 7]. Other methods rely on integrating diverse high-throughput data, such as gene coexpression or protein complex co-purification, which limits their use to well-studied organisms with extensive experimental datasets [4]. Machine learning, particularly deep learning approaches such as graph-based learning [8] and other advanced frameworks [9] applied to sequence or structural data [6], has shown great promise [7]. However, these methods typically require large, well-annotated training sets specific to particular organisms or interaction types, which may not always be available for ab initio discovery in less-characterized species. Algorithms designed to predict protein-protein interactions can utilize Gene Ontology [2] or combine it with other strategies [10]. However, we have shown that more than just Gene Ontology is needed to produce satisfactory results in the prediction process [11]. We extensively reviewed the literature considering recent years, and no state-of-the-art software can predict interaction networks using exclusively genomic data efficiently.

The main contributions of this work are thus: (i) An alignment-free protein similarity prediction method integrated into GenPPi, utilizing a Random Forest classifier trained on biophysical features to identify similarities below conventional sequence identity thresholds effectively. (ii) The novel Reduced Interaction Sampling (RIS) algorithm is designed to manage the computational complexity arising from increased interaction candidates in sensitive prediction modes, thereby enabling scalable analysis. (iii) Rigorous validation of the machine learning model’s performance across diverse bacterial genomes and a comprehensive evaluation of the RIS algorithm’s impact on network topology preservation. (iv) An enhanced, parallelized, and opensource GenPPi 1.5 tool that significantly advances the state-of-the-art for practical *ab initio* PPI network prediction, particularly for understudied genomes. Moreover, the possibility of crafting a training model empowers researchers to optimize GenPPi’s performance for their specific organisms of interest, potentially achieving even greater accuracy and relevance in their PPI predictions. GenPPi thus aims to fill a critical gap by providing a robust platform for de novo network generation, especially for organisms outside the scope of exhaustive experimental study, complementing existing databases and structure-dependent predictive tools.

This paper elaborates on these contributions, presenting the methods employed for model training and RIS development (Section 2), evaluating their performance and impact through extensive testing (Section 3), and discussing the results in the context of current PPI prediction challenges (Section 4).

### 1.1 State of the Art

In the context of predictions of protein interactions, the STRING (Search Tool for the Retrieval of Interacting Genes/Proteins) stands out as one of the leading stateof-the-art tools. STRING is a database that integrates known and predicted protein interactions, encompassing multiple sources of evidence, including biochemical experiments, genetic associations, and gene coexpression. Its applicability extends to more than 2000 organisms, allowing comparative analyses between species. The platform offers an intuitive interface for visualizing interaction networks, facilitating the identification of functional modules and the interpretation of complex biological contexts.

The constant updating and inclusion of new functionalities make STRING a valuable tool for bioinformatics and systems biology researchers, significantly advancing knowledge about protein interactions and their implications in health and disease [12]. Other tools are available for predicting protein-protein interaction networks, but not all are applicable or comparable to GenPPi in all contexts. For example, some of these tools address specific organisms, do not allow data upload for analysis, or are limited to data from multicellular organisms (eukaryotes). These limitations make it impractical to compare these tools directly with GenPPi, especially concerning unpublished, little-known, or understudied complete genomes. GenPPi, on the other hand, allows the prediction of interaction networks from any genome of bacteria or archaea. The flexibility in creating interaction networks distinguishes GenPPi from other methodologies, making it particularly useful for researchers working with genomes that other tools may not be able to cover. Therefore, while being aware of the other tools and methodologies available is helpful, it is crucial to understand the limitations that prevent a direct comparison with GenPPi.

For example, in [10], which utilizes information from known complexes, gene expression profiles, Gene Ontology terms, and subcellular localization information to discover essential proteins, the algorithm is theoretical. It lacks an associated software for direct implementation in the GenPPi code.

Regarding [13], which proposes a new method to predict unknown protein-protein interactions, it is important to clarify that the associated software presented functionality problems despite its theoretically interesting proposal. Attempts to execute the code made available by the authors were unsuccessful, and our requests for assistance still need to be answered. Therefore, it was not possible to compare our results with theirs.

In addition, during the development of GenPPi, it was demonstrated that the interaction network accuracy produced using Gene Ontology was inferior to an *ab initio* generation [11].

The topological analysis of protein-protein interaction networks (PPI) is a relevant aspect of bioinformatics and systems biology, serving as one of the primary motivations for developing version 1.5 of GenPPi. A study entitled *Topological scoring of protein interaction networks* [14] reinforces the need for tools such as GenPPi, which facilitate topological analysis in PPI networks, contributing to a better understanding of protein interactions and their functional implications.

## 2 Methods

### 2.1 Features

The source code, named *Features.lisp*, is a recent addition to GenPPi in version 1.5, an in-house program that utilizes a file called *propensity.dat* as a basis for mapping specific protein features. This file contains specific values for each amino acid, allowing the program to create a propensity index for each characteristic of each amino acid present in all proteins in a multi-fast file. The data in the *propensity.dat* file represent different physicochemical characteristics of the amino acids, such as acidity/basicity (BASIC, ACID), polarity (POLAR, NONPOLAR), mass (MASS, MASSMR) and other specific indices (PARJ860101, JOND750101, EISD840101, JURD980101), in a total of ten characteristics available in the database of the AAindex website [15].

Each row of the *propensity.dat* file represents a characteristic, and each column is an amino acid. The numerical values represent the propensity of each amino acid for this characteristic. These indices were selected based on their effectiveness in classifying proteins according to the most common subcellular sites in prokaryotic organisms, such as the membrane, cytoplasm, cell wall surface, and secreted proteins[16].

The program reads the FASTA file, which contains several amino acid sequences, and the propensity file, which contains information about the physicochemical properties of amino acids. For each sequence in the FASTA file, the *Features* program calculates a histogram that counts the occurrence of each amino acid in the sequence. In addition, it calculates the weighted sum of the physicochemical properties for the entire sequence and the start, end, and middle regions of a sequence.

For each trait, the *Features* program calculates the value along the entire protein. Additionally, it segments the protein into three parts: the beginning, middle, and end, resulting in four values for each trait. This results in a subtotal of forty values per protein. In addition, simply counting the amount of each amino acid adds twenty more numerical characteristics to each protein, totaling sixty values defining a protein’s propensity index. A more significant number of discriminative features allows for a more granular analysis of protein properties, as different regions of a protein can have different amino acid compositions and, therefore, different physicochemical properties.

### 2.2 Random Forest

The incorporation of Random Forest as a machine learning algorithm in GenPPi presents a solution to enhance the ability to predict protein-protein interaction networks (PPIs) in bacterial genomes, a limitation faced by GenPPi 1.0, which resulted in non-optimal sampling of similar proteins. The challenge faced was to identify interactions between proteins with levels below 90%. Exact alignment or heuristic algorithms, such as blast, become computationally costly, especially in complex pangenomes, which is why we have sought alternative ways to identify protein similarity.

The choice of the Random Forest algorithm as a solution to the problem is justified by its ability to efficiently handle complex and high-dimensional datasets, as is the case with the genomic and proteomic information involved in the GenPPi[17]. Random Forest builds a collection of decision trees, each trained on a random subset of the available data and variables. This approach promotes diversity among trees, preventing dominant traits from exerting an excessive influence on the model and contributing to better generalization. In this work, we aimed to find a balance, improving sensitivity for lower-similarity pairs without generating an intractably large number of candidates for the downstream genomic analyses. Each tree forms a structure similar to a flowchart, contributing to joint decision-making [18].

We based the choice of the *cl-random-forest* library for incorporation into GenPPi on the need for robust and efficient implementation of Random Forest in Common Lisp [19]. The cl-random-forest library is an implementation of the Random Forest algorithm for SBCL. This library supports multiclass classification and univariate regression and is notable for its efficiency and accuracy, outperforming other popular implementations such as scikit-learn (Python/Cython) and ranger (R/C++). In addition, *cl-random-forest* includes a global refinement of Random Forest, as described in [20], making it even faster and more accurate than the standard implementation. This choice also facilitated integration within GenPPi’s existing Common Lisp codebase.

The code used to train the Random Forest model used in GenPPi 1.5 is available in the GitHub repository https://github.com/santosardr/non-CSPs/tree/main/src/lisp. This file depends on the *cl-random-forest* and *cl-store* libraries installed via Quicklisp. Different training and testing sets were used with this interface to generate the final model for predicting similarities between protein pairs used in this new version of GenPPi.

### 2.3 Database

We compared the results obtained by the GenPPi software with the STRING data. STRING is recognized as a gold standard in protein interaction analysis, providing information on known interactions and predictions of interactions between proteins. For this purpose, we used the PostgreSQL relational database management system.

The database used for this analysis is available in the GenPPi repository on GitHub, under the name *compareba.dump*, since the study organism for comparison is *Buchnera aphidicola*. We designed the database *Ba ak* for genome-related analyses, which is used as an example in this study. It consists of two main tables: *protein* and *related*. In Table *protein*, we stored the identifiers of the proteins and from which genome they originate, obtained from both the STRING interaction network and the GenPPi interaction network. The *related* table stores the interactions between proteins of the same genome, allowing the comparison of interactions between proteins.

We developed the *count intersect distinct* function to analyze protein interactions accurately. This function returns a table with the name of the genome and the number of unique interactions performed, considering the interactions with and without repetitions. In the context of STRING, where the interaction between A and B is considered different from that between B and A, the count of interactions without repetitions equals half the total interaction count. On the other hand, in the GenPPi, repeated interactions are removed using the Gephi program, which is commonly employed in this work to visualize the interaction networks generated by the GenPPi [21].

### 2.4 Experiments

#### 2.4.1 Genomes Training and Testing

We turned to the National Center for Biotechnology Information (NCBI) due to its vast database and the availability of genomes of model pathogenic bacteria. We used the genomes of several species, including *Aeromonas hydrophila, Bacillus anthracis, Bacteroides fragilis, Borrelia burgdorferi, Clostridium botulinum, Clostridium tetani, Corynebacterium Jeikeium, Enterobacter huaxiensis, Enterococcus faecalis, Escherichia coli, Francisella tularensis, Helicobacter pylori, Moraxella Catarrhalis, Morganella morganii, Mycobacterium avium, Mycobacterium tuberculosis, Providencia rettgeri, Pseudomonas aeruginosa, Salmonella enterica, Staphylococcus aureus, Streptococcus pyogenes, Vibrio cholerae and Yersinia pestis*. We based the selection of these species on their relevance as pathogenic agents and on their extensive study in the scientific literature.

The roles of these genomes in our study were distinct and carefully separated to ensure the robust development and evaluation of our model. Specifically: (i) Training Set Generation: Nine genomes (detailed in Section 2.4.2) were used exclusively for generating the primary training dataset for the Random Forest (RF) model. (ii) RF Hyperparameter Optimization Test Set: A completely independent test set was constructed for the crucial step of RF hyperparameter selection and optimization. This set was derived from the combined proteomes of *Corynebacterium pseudotuberculosis* (2033 proteins) and *Corynebacterium glutamicum* (2970 proteins). An all-versus-all BLASTP comparison of these 5003 proteins yielded 837 positive pairs (defined by *>*65% sequence identity), forming the basis for evaluating different RF training configurations (see Section 2.4.2 and Table 1). (iii) General RF Model Evaluation: A broader set of 26 diverse genomic test sets, including both intra- and inter-genome comparisons (as listed in Table 2), was used to assess the performance of the finalized RF model. (iv) Benchmarking and RIS Evaluation: Finally, the *Buchnera aphidicola* (Ba ak) genome was employed for the detailed evaluation of the RIS algorithm (Section 2.6) and for the comparative benchmarking against GenPPi 1.0 and the STRING database (Section 3.5). Critically, the Ba ak genome was not used during any phase of RF model training or its hyperparameter tuning.

**Table 1.** Optimization of Random Forest Hyperparameters for Protein Similarity Classification. This table details the performance (Sensitivity, Specificity, F1-Score) of Random Forest models trained with standard library defaults versus empirically chosen custom hyperparameters. Models were evaluated across seven different negative-to-positive instance ratios in the training set, with all performance metrics reported on the independent *C. pseudotuberculosis* + *C. glutamicum* test set to guide final model selection.

| Training Set<br>(Approx. Ratio)<br>(Neg:Pos) | Standard Parameters |  |  | Custom Parameters |  |  |
| --- | --- | --- | --- | --- | --- | --- |
|  | Sensitivity<br>(Test Set) | Specificity<br>(Test Set) | F1-Score<br>(Test Set) | Sensitivity<br>(Test Set) | Specificity<br>(Test Set) | F1-Score<br>(Test Set) |
| 1 (1:1) | 0.4803 | 0.8712 | 0.5160 | 0.9988 | 0.3868 | 0.5238 |
| 2 (2:1) | 0.2342 | 0.9762 | 0.3590 | 0.9534 | 0.7235 | 0.6879 |
| 3 (**3:1**) | 0.2055 | 0.9814 | 0.3261 | **0.8829** | **0.8466** | **0.7556** |
| 4 (4:1) | 0.1649 | 0.9851 | 0.2727 | 0.6428 | 0.9370 | 0.7028 |
| 5 (5:1) | 0.1243 | 0.9895 | 0.2151 | 0.5651 | 0.9471 | 0.6565 |
| 6 (6:1) | 0.1075 | 0.9923 | 0.1903 | 0.5197 | 0.9536 | 0.6273 |
| 7 (7:1) | 0.0980 | 0.9948 | 0.1760 | 0.4659 | 0.9576 | 0.5856 |

Test set (*C. pseudotuberculosis* + *C. glutamicum*): 2477 Negative, 837 Positive instances. Custom Parameters: :n-tree 500, :max-depth 30, :n-trial 10.
Standard parameters are the defaults provided by the cl-random-forest library.

**Table 2.** Performance of the Finalized GenPPi Random Forest Model Across Diverse Bacterial Genome Test Sets. This table presents key classification metrics (True Negatives, False Negatives, True Positives, False Positives, Sensitivity, Specificity, and F1-Score) for the optimized Random Forest model (trained with a 3:1 negative-to-positive ratio and custom hyperparameters). Performance was assessed on 26 different intraand inter-genome test sets to evaluate its generalization capability for identifying protein similarity.

| Genomes /<br>Test Set | Confusion Matrix Counts |  |  |  | Performance Metrics |  |  |
| --- | --- | --- | --- | --- | --- | --- | --- |
|  | TN | FN | TP | FP | Sensi-<br>tivity | Speci-<br>ficity | F1-Score |
| *Aeromonas hydrophila* | 1713 | 14 | 26 | 35 | 0.6500 | 0.9799 | 0.5149 |
| *Aeromonas hydrophila* - *Bacteroides fragilis* | 3548 | 112 | 84 | 273 | 0.4286 | 0.9286 | 0.3038 |
| *Aeromonas hydrophila* - *Clostridium botulinum* | 3347 | 61 | 68 | 93 | 0.5271 | 0.9730 | 0.4690 |
| *Aeromonas hydrophila* - *Enterobacter huaxiensis* | 4277 | 39 | 69 | 112 | 0.6389 | 0.9745 | 0.4775 |
| *Bacteroides fragilis* | 1835 | 98 | 58 | 238 | 0.3718 | 0.8852 | 0.2566 |
| *Bacteroides fragilis* - *Clostridium botulinum* | 3469 | 145 | 100 | 296 | 0.4082 | 0.9214 | 0.3120 |
| *Clostridium botulinum* | 1634 | 47 | 42 | 58 | 0.4719 | 0.9657 | 0.4444 |
| *Enterobacter huaxiensis* | 2564 | 25 | 43 | 77 | 0.6324 | 0.9708 | 0.4574 |
| *Enterobacter huaxiensis* - *Clostridium botulinum* | 4198 | 72 | 85 | 135 | 0.5414 | 0.9688 | 0.4509 |
| *Enterobacter huaxiensis* - *Bacteroides fragilis* | 4399 | 123 | 101 | 315 | 0.4509 | 0.9332 | 0.3156 |
| *Morganella morganii* | 938 | 18 | 95 | 91 | 0.8407 | 0.9116 | 0.6355 |
| *Providencia rettgeri* | 1066 | 62 | 207 | 125 | 0.7695 | 0.8950 | 0.6889 |
| *Yersinia pestis* | 1625 | 1461 | 3768 | 51 | 0.7206 | 0.9696 | 0.8329 |
| *Streptococcus pyogenes* | 320 | 9 | 13 | 7 | 0.5909 | 0.9786 | 0.6190 |
| *Mycobacterium avium* | 2463 | 164 | 266 | 493 | 0.6186 | 0.8332 | 0.4474 |
| *Mycobacterium avium* - *Corynebacterium jeikeium* | 2843 | 270 | 392 | 543 | 0.5921 | 0.8396 | 0.4909 |
| *Mycobacterium avium* - *Moraxella catarrhalis* | 2593 | 173 | 285 | 506 | 0.6223 | 0.8367 | 0.4564 |
| *Corynebacterium jeikeium* | 380 | 106 | 126 | 50 | 0.5431 | 0.8837 | 0.6176 |
| *Corynebacterium jeikeium* - *Moraxella catarrhalis* | 510 | 115 | 145 | 63 | 0.5577 | 0.8901 | 0.6197 |
| *Moraxella catarrhalis* | 130 | 9 | 19 | 13 | 0.6786 | 0.9091 | 0.6333 |
| *Helicobacter pylori* | 171 | 21 | 21 | 6 | 0.5000 | 0.9661 | 0.6087 |
| *Helicobacter pylori* - *Borrelia burgdorferi* | 284 | 331 | 449 | 73 | 0.5756 | 0.7955 | 0.6897 |
| *Helicobacter pylori* - *Pseudomonas aeruginosa* | 3570 | 81 | 82 | 175 | 0.5031 | 0.9533 | 0.3905 |
| *Pseudomonas aeruginosa* | 3399 | 60 | 61 | 169 | 0.5041 | 0.9526 | 0.3476 |
| *Pseudomonas aeruginosa* - *Borrelia burgdorferi* | 3512 | 370 | 489 | 236 | 0.5693 | 0.9370 | 0.6174 |
| *Borrelia burgdorferi* | 113 | 310 | 428 | 67 | 0.5799 | 0.6278 | 0.6942 |
| **Mean** | **-** |  |  |  | **0.5734** | **0.8998** | **0.5136** |

Data from model organisms, such as those found in the NCBI, can provide a solid basis for future investigations, allowing the confirmation of hypotheses and conclusions reached in our work. Additionally, the availability of these genomes enables the creation of representative datasets and the performance of comparative analyses, thereby enhancing the robustness and validity of the results obtained in our experiments.

#### 2.4.2 Generating the Machine Learning Model

In the creation of the training file, we selected proteins from nine medically important genomes: *Bacillus anthracis, Clostridium tetani, Enterococcus faecalis, Escherichia coli, Francisella tularensis, Mycobacterium tuberculosis, Salmonella enterica, Staphylococcus aureus, and Vibrio cholerae*. The *src/tools/makearff.bash* script performs a series of manipulations on text files containing organic information, applying filters and converting them to the ARFF format, which is essential for training the algorithm. We used *src/tools/makearff.bash* to compare the nine genomes. This process generated a file with proteins identified with their characteristics and the class to which they belong (positive or negative).

The original full training dataset exhibited a significant imbalance between positive (8,858) and negative (318,309) protein pairs. Recognizing that such class imbalance can negatively impact classifier performance, we compared the performance of the Random Forest classifier using the default parameters provided by the cl-random-forest library against a set of empirically chosen custom parameters. Crucially, this hyperparameter optimization, including the determination of the optimal negative-to-positive instance ratio for the training data and the selection of custom classifier settings, was conducted exclusively using the independent test set derived from the combined *C. pseudotuberculosis* and *C. glutamicum* proteomes (837 positive pairs, with negative pairs drawn from the same combined set but lacking *>*65% identity). This *Corynebacterium* test set was not part of the RF training data nor the final Ba ak benchmarking dataset. This comparison was conducted across seven training subsets with varying negative-to-positive instance ratios (from 1:1 to 7:1), all evaluated on the independent *Corynebacterium* test set (Table 1).

**Algorithm 1.**
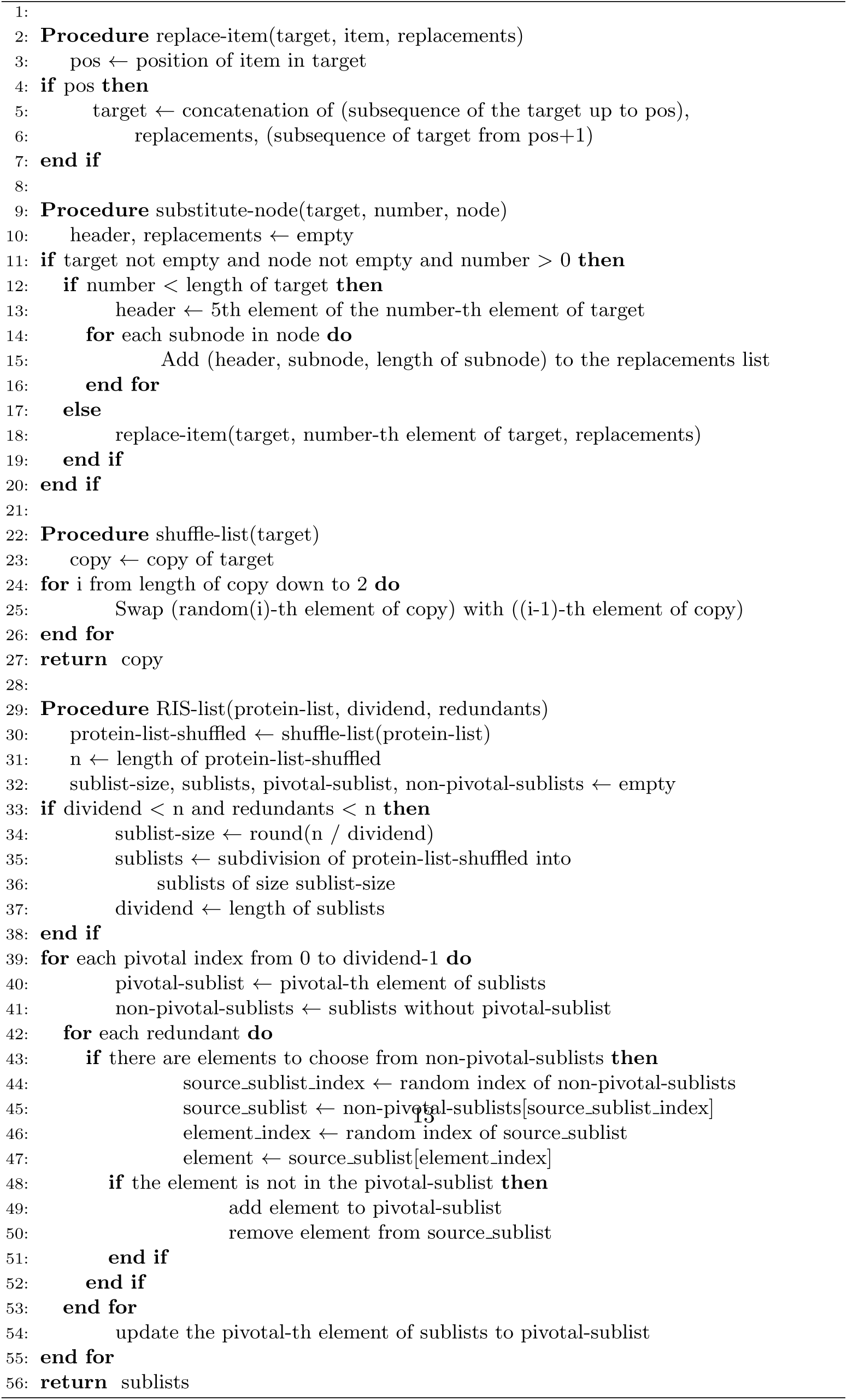
The Reduced Interaction Sampling (RIS) Algorithm. This pseudocode outlines the RIS procedure designed to manage computational complexity in PPI network generation. RIS stochastically samples interactions from large phylogenetic profiles by first shuffling the proteins within a profile and then subdividing the list into smaller, manageable sublists. Interactions are primarily generated within these sublists, with a mechanism to ensure some overlap and maintain network connectivity, thereby reducing-edge density while preserving core topological features.

The results in Table 1 highlight a clear trade-off. The standard parameters consistently yielded very high Specificity, particularly as the negative:positive ratio increased, but suffered from drastically reduced Sensitivity (Recall), rendering them unsuitable for GenPPi’s discovery-oriented goals as they would miss most true similarities at higher, more realistic imbalance ratios.

Conversely, the custom parameters demonstrated significantly higher Sensitivity across all tested ratios. Given that GenPPi’s primary goal is not only to confirm known interactions but also to facilitate the discovery of potentially novel ones, which often depend on identifying less obvious similarities missed by high-stringency methods, maximizing Sensitivity without excessively compromising Specificity was the primary selection criterion. The F1-score for the custom parameters peaked at the 3:1 ratio when evaluated on this specific *Corynebacterium* test set, offering a strong balance between Sensitivity (0.8829) and Specificity (0.8466). While the 1:1 and 2:1 ratios offered even higher sensitivity, their specificity was considerably lower, leading to potentially unmanageable false positives. The subsequent filtering steps within GenPPi, which rely on analyzing genomic context, are designed to handle a moderate level of false positives. However, starting with a reasonable specificity is still beneficial. Therefore, based solely on this evaluation on the independent *Corynebacterium* (*pseudotuberculosis* and *glutamicum*) test set, we chose the 3:1 ratio trained with custom parameters for the final model. It provides the best input for the overall process by maximizing the capture of true potential similarities for downstream validation while maintaining acceptable specificity, representing the optimal compromise for GenPPi’s objectives.

The custom hyperparameters used for the final model, justified by this comparative analysis, which shows their advantage over library defaults for our specific task, were configured as follows. The final model utilized 500 decision trees (:n-tree 500). To prevent overfitting while allowing sufficient complexity, the maximum depth of each tree (:max-depth) was programmatically set to half the number of input features plus one (resulting in a depth of 30 for our 60 features). The number of random feature subsets evaluated at each node split (:n-trial) was increased to 10 from the default of 1 to improve split quality potentially. We used default values for bagging ratio (:bagging-ratio 1.0) and minimum samples per leaf node (:min-region-samples 1).

A key strength of GenPPi lies in its flexibility regarding the underlying machine learning model. While we provide a pre-trained model optimized using the 3:1 ratio and the custom parameters described above, users are not limited to this default. The GenPPi architecture enables the creation of custom models tailored to specific taxonomic groups or research questions. To facilitate this, we provide the maker script (src/tools/makearff.bash, available on the GenPPi GitHub repository), which converts protein sequence data in FASTA format into the ARFF format required for training the Random Forest classifier. Users can then follow the instructions provided in the repository (src/tools/crafting-RF-training-model.lisp) to generate a binary model file (model.dat) using any training data and desired parameters. Users should know the need to install the library cl-random-forest before crafting their training models.

We can call the program with the following command: *./randomforest similar test.arff xxxx similar training.arff*, where the second parameter *(similar test.arff)* is the test file containing the target genome proteins, the third parameter *(xxxx)* indicates the number of negative proteins present in the test file, in digit format. It is essential to note that we have carefully structured the ARFF file to ensure that all negative proteins precede positive ones. This ordering is critical, as we use true labels for each data instance in training. The Random Forest program, implemented in Lisp, utilizes this information to calculate the model’s accuracy as it is being constructed, taking into account both successes and errors. The last parameter represents the training file. Like the test file, the training file follows the pattern of negative instances preceding positive ones. This compliance enables the interchangeability of training and test files, providing flexibility in evaluating and validating machine learning models.

#### 2.4.3 Using the GenPPi-trained model

The GenPPi initially employed the simple counting of the occurrences of the twentysix amino acids (including six non-standardized) as a method of analysis. However, in the enhanced version, incorporated by the sixty features generated by the *Features* program, in the default mode of execution, called *Features* mode, we consider two proteins similar if they present at least twenty-five of these characteristics with a maximum difference of one or two units. This last number is a value obtained empirically. Changing this number affects the ability to generate inferences of correct similarities. If this number were increased too much, for example, close to the sixty features that GenPPi 1.5 uses, GenPPi 1.5 in *Features* mode would pass like GenPPi 1.0 when it used twenty-six amino acid possibilities.

We employed the same sixty features used by the *Features* algorithm in the *clrandom-forest* program, which performs machine learning to create more complex networks of proteins. We can trigger the GenPPi machine learning mode through the optional *-ml* parameter. To use the machine learning mode, the user must download the previously trained Random Forest model to recognize similar protein pairs with more than 65% identity and size alignments as determined by BLASTP [22].

In the new version of GenPPi 1.5, we standardized some parameters to simplify the user experience. The *-ppcomplete* parameter, used to perform PPI predictions by the conserved phylogenetic profiling method without applying filters, is now enabled by default; the program will make the predictions without reducing the number of predicted interactions.

In addition, the type of neighborhood expansion of a recurrent gene controlled by the *-expt* parameter is now *fixed*. The size of the main window *(-w1)* and the required amount of genes conserved within window 1 *(-cw1)*, which were set to ten and four, respectively, are now also default values. Therefore, without extra parameters, GenPPi 1.5 will run with these default settings, simplifying the process and allowing users to focus more on interpreting the generated results.

Once one clones our example repository; it is possible to perform tests on the *Buchnera Aphidicola* genome. The program can be run with the following command (in the Linux operating system): *./genppi -dir test1*. The first parameter represents the version of the GenPPi program selected for the machine, with multiple builds available for different memory usage limits. The second parameter *(’-dir’)* indicates the name of the directory where we stored the genomes in the form of multifasta files (one file per genome), and the last parameter *(’test1’)* is the name of the directory in question. For instance, we can fire the program’s machine learning mode (in Linux) with the command: *./genppi -dir test1 -ml*. The parameters are the same, except the last one *(’-ml’)*, which instructs the program to use machine learning to identify similar proteins.

### 2.5 RIS (Reduced Interaction Sampling) Algorithm

When employing machine learning for PPI prediction in GenPPi, a significant increase in identified interactions was observed compared to approaches that do not utilize machine learning. This increase, often driven by large conserved phylogenetic profiles involving hundreds of genes, represents an additional challenge for the subsequent topological analysis of the resulting network due to combinatorial explosion. For instance, a profile with *x* = 500 conserved genes could potentially generate 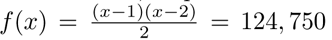 edges, rapidly leading to computationally intractable networks. This computational bottleneck highlights why simply maximizing the recall of any similarity classifier is insufficient; a reduction strategy becomes essential.

To mitigate this complexity while retaining the sensitivity gained from machine learning, the RIS (Reduced Interaction Sampling) algorithm, previously referred to as ’split,’ was developed and incorporated into the GenPPi 1.5 source code (Algorithm 1). The core idea is to sample interactions from large phylogenetic profiles rather than generating all possible pairs, thereby reducing the edge density without drastically altering the network’s core structure. RIS operates through three main steps:

1. **Shuffle:** The list of proteins within a large phylogenetic profile is randomly shuffled to eliminate any ordering bias.
2. **Subdivide:** The shuffled list is divided into multiple smaller sublists. We typically set the number of sublists to 90% of the original profile size.
3. **Independent Processing:** Each sublist is treated as an independent, smaller phylogenetic profile, and we generate interactions only *within* each sublist. A mechanism also ensures some overlap by randomly distributing several proteins among the sublists to maintain connectivity.

This strategy significantly reduces the number of pairwise comparisons and resulting edges, making downstream network analysis feasible even for complex pangenomes. The crucial question in the following sections is whether this sampling process preserves the network’s essential topological features.

### 2.6 RIS Evaluation Methodology

To rigorously evaluate the impact of RIS on network topology, we conducted extensive computational experiments using the *Buchnera aphidicola* (Ba ak) genome as a reference. We generated 50 independent network replicates for each of GenPPi’s operational modes: *Features* (relying on biochemical characteristics) and *Machine Learning* (incorporating the Random Forest classifier). This replication strategy aimed to account for the stochasticity introduced by RIS’s sampling process.

The experimental design involved varying key parameters to assess their influence on network stability:

- **Operational Mode**: Features vs. Machine Learning.
- **Number of Input Files (**NrFiles): Ranging from 10 to 50, simulating different dataset sizes.
- **Top N Nodes**: Selecting the top N nodes based on centrality, ranging from 10 (approx. 1.8% of the proteome) to 200 (approx. 35.8%), to evaluate RIS’s effect at different levels of network granularity.
- **Topological Metrics**: Focusing on Degree, Betweenness Centrality, and Bridging Centrality as criteria for selecting Top N nodes.

Our experimental design resulted in 180 distinct experimental configurations. For each generated network (in DOT format), we extracted topological metrics using Gephi [21] and saved them into CSV files for comparative analysis.

To quantify the stability and structural preservation under RIS, we employed a suite of statistical metrics comparing the replicates within each configuration:

- **Topological Metrics**: Degree (identifying hubs), Betweenness Centrality (identifying bottlenecks), and Bridging Centrality (identifying connectors between communities) [23]. We used these as criteria for Top N selection and as measures of structural change.
- **Weighted Global Presence Mean (MGP)**: A custom metric developed to measure the consistent presence of critical nodes across replicates, weighted by their frequency of appearance, mitigating bias from sparsely appearing nodes.
- **Jaccard Coefficient**: To quantify the similarity between the sets of Top N nodes across pairs of replicates [24].
- **Fleiss’ Kappa (Adapted)**: To assess the multi-rater agreement (concordance) among all replicates on the selection of Top N nodes, correcting for chance agreement [25].
- **Kendall Tau Rank Correlation Coefficient**: To measure the consistency of the hierarchical ranking of the Top N nodes between pairs of replicates [26]. We specifically focused on the Kendall Ratio Positive (proportion of concordant pairs).
- **Kolmogorov-Smirnov (KS) Test (Multi-Sample)**: To compare the empirical distribution functions (CDFs) of the chosen topological metric (e.g., Degree) across all replicates within a configuration, assessing distributional similarity [27].

These metrics collectively allowed for a multifaceted assessment of whether RIS maintains the predicted PPI networks’ global and local structural properties, hierarchical organization, and distributional characteristics.

### 2.7 Parallelism

The first GenPPi version was monoprocessed. We implemented parallelism in three code bottlenecks. The first critical point was the call of the function *call-similar-test* to determine a high similarity between protein pairs, the function now responsible for calling our Machine Learning implementation. We can find the second crucial point in the functions prefixed by *phylogenetic-profiles*responsible for inferring a conserved PP under different conditions triggered by specific parameters. We can find the last point at the functions *execute-expansion-fixed* and *execute-expansion-dynamic* performing the conserved neighborhood checking among proteins. The GenPPi will utilize all the processors available in a computer except for two, allowing for better machine control. When we analyze two or more genomes whose organisms are evolutionarily close, the tendency is for the number of genes in conserved phylogenetic profiles to increase significantly. For example, it is very common to have phylogenetic profiles conserved with more than five hundred genes in organisms with more than two thousand genes. Therefore, the number of edges initially created would be 124,251 edges, with each gene having 499 edges connected to the others. For another with 700 conserved genes, we would have 243,951 edges, with each gene having 699 edges. With a few large profiles conserved in a genome, it is very common that at the end of the generation of the interaction network, we have edges that exceed half a million or even a million. The final number of edges is proportional to the level of similarity between the genomes. To slow down this rapid growth in the number of edges and average degrees of the graph’s vertices, we thought of a process that could make a meaningful sampling of the interactions existing in phylogenetic profiles with several hundred conserved genes. Sampling aims to create as many interactions as possible between the genes, with many genes, without necessarily including all the edges. In this way, we were able to show that gene *A* interacts with gene *B* without necessarily including all the interactions of genes *A* and *B*. We refer to this sampling approach as RIS, alluding to how it works by dividing a profile’s large set of genes into many smaller portions. The RIS generates several smaller profiles, each equal to 90% of the size of the original profile, and places in each derived profile a smaller number of randomly selected genes in each derived profile.

Initially, the *RIS* promotes a disorder of the list of similar proteins known as *shuffle*. Later, we subdivided this list into several sublists equal to 90% of the number of elements in the original list. We randomly assign each sublist about half minus one element of the undivided unordered list. The program is then run separately on each sub-list, reducing the number of possible combinations and, consequently, the total number of edges in the network. Splitting and executing separately into sub-groups effectively reduces the number of interactions, facilitating topological analysis of the resulting network.

When considering this strategy, a relevant question is the fate of proteins that do not form edges. Although we miss some interactions, the tests carried out showed that after the application of the *RIS*, the result of the network of interactions practically did not change in terms of analyses of the topological importance of nodes, that is, in addition to reducing the number of edges in the network, which facilitates the analysis of the network of protein-protein interactions (PPI), The quality of the underlying data was not compromised, maintaining node differentiation in centrality measures. The explanation is that the decrease in the degree of the nodes was done homogeneously due to randomness, preserving the ranking of the most connected nodes. This approach provides an effective solution to address the increased number of interactions resulting from machine learning while maintaining the quality and integrity of the data for subsequent analysis by the PPI network.

The application of these methods yielded the results presented below.

## 3 Results

### 3.1 Performance Analysis of the Random Forest Classification Model

The Random Forest (RF) model was configured using hyperparameters optimized as described in Section 2.4.2, based on evaluations against an independent test set derived from *C. pseudotuberculosis* and *C. glutamicum* (see Table 1 in Section 2.4.2 for details of this optimization). Table 2 now presents the performance metrics of this finalized RF model, evaluated across 26 diverse genome test sets. Test data for each genome or pair were created by identifying protein pairs with *>*65% sequence identity via BLASTP (positive class) versus those below this threshold (negative class), following the methodology described earlier.

The evaluation across these independent test sets reveals the performance profile of the optimized model. The average Specificity is at approximately 90.0%, while the average Sensitivity is 57.3%. Consequently, the average F1-Score, which balances precision and recall (Sensitivity), is approximately 51.4% (Table 2, last row).

The evaluation across these independent test sets reveals the performance profile of the optimized model. The average Specificity is at approximately 90.0%, while the average Sensitivity is 57.3%. Consequently, the average F1-Score, which balances precision and recall (Sensitivity), is approximately 51.4% (Table 2, last row).

This performance profile, characterized by high Specificity and moderate Sensitivity, warrants careful interpretation within the context of the overall GenPPi pipeline. While moderate sensitivity might seem disadvantageous in isolation, high specificity is crucial for GenPPi’s operational efficiency. The downstream analyses within GenPPi operate on the pairs initially flagged as similar by the RF model. Given that the potential search space for pairs is vast (O(N^2) for N proteins), minimizing False Positives (FP) through high Specificity is paramount. A model with significantly lower specificity, even if slightly more sensitive, would generate an unmanageably large number of false leads, rendering the subsequent computationally intensive genomic context analyses intractable.

Furthermore, maximizing sensitivity in the initial similarity screen is balanced against the need for manageable downstream processing. GenPPi’s core goal is to infer interaction networks based on conserved *groups* of genes. Identifying a sufficient number of correct similarity links *within* such groups, enabled by the model’s moderate sensitivity (0.5734 average), is often enough to flag the entire set of proteins for interaction inference based on genomic evidence. Discovering *every* possible similar pair might be redundant.

Significant variability in performance across different test sets is still observed (Table 2). For example, the model achieves high Sensitivity and F1-Score for organisms like *Morganella morganii* (Sens=0.841, F1=0.635) and *Providencia rettgeri* (Sens=0.770, F1=0.689), and particularly high precision for *Yersinia pestis* contributing to a strong F1-score despite moderate sensitivity (Sens=0.721, F1=0.833). Conversely, sensitivity and F1-score remain lower for pairs involving more distant organisms or those with potentially unique protein characteristics, such as *Pseudomonas aeruginosa* pairs (e.g., Sens=0.504, F1=0.348 for intra-genome test) or some inter-genome tests involving *Bacteroides fragilis* (e.g., Sens=0.408, F1=0.312 vs *C. botulinum*), while still generally maintaining high Specificity. The variability of RF results highlights the inherent challenges in creating a universally performing similarity model based solely on these features. It underscores the importance of evaluating performance across diverse genomic backgrounds. It is important to note that users can further optimize performance by training custom models using their datasets, as described in SubSection 2.4.2.

Therefore, the trained model strikes a balance, prioritizing the confident rejection of non-similar pairs (high Specificity) while capturing a significant portion of potentially similar pairs (moderate Sensitivity). The resulting F1-Score reflects this balance, deemed optimal for GenPPi’s specific architecture, where the initial RF classification acts as a critical filter to provide a tractable, high-confidence set of candidate pairs for the more biologically informative downstream genomic analyses. The objective is not merely to replicate BLAST results at a lower threshold but to efficiently identify plausible candidates for interaction based on combined evidence.

### 3.2 RIS Evaluation Results

The comprehensive evaluation across 180 configurations yielded extensive data, publicly available as detailed in [28]. Here, we summarize the key findings regarding the impact of the RIS algorithm.

**Features Mode:** In this mode, the RIS algorithm demonstrated a negligible impact on network topology. Across nearly all configurations (85 out of 90), the Weighted Global Presence Mean (MGP) was 100%, with the lowest value observed being 99.76%. Similarly, Jaccard coefficients were consistently near 1.0, and Fleiss’ Kappa indicated perfect agreement. These metrics outcomes demonstrate that when using the deterministic Features mode, RIS effectively reduces computational load without compromising the structural stability or reproducibility of the network predictions.

**Machine Learning Mode:** As anticipated, introducing RIS in the Machine Learning mode led to greater variability than the Features mode, primarily due to the stochastic nature of both the ML predictions and the RIS sampling. However, the analysis revealed clear trends related to the Top N parameter:

- **Impact of** Top N **on Stability:** The Top N parameter emerged as the most critical factor influencing network stability. Increasing Top N consistently led to higher stability across all metrics. For instance, MGP values (averaged across metrics and FilesNum) increased substantially, ranging from approximately 13-33% for Top N=10 to 70-76% for Top N=200 (Fig. 1). The results from Fig. 1 indicate that analyzing a larger fraction of the network’s core structure mitigates the random effects introduced by RIS.
- **Preservation of Node Hierarchy:** The Kendall Tau correlation analysis showed remarkable stability in ranking critical nodes. For Top N *≥* 100, the Kendall Ratio Positive approached or reached 1.0 across most configurations (Fig. 1), indicating preservation of the relative importance hierarchy of the main nodes despite the edge reduction performed by RIS.
- **Distributional Similarity:** The multi-sample KS test distance generally decreased as Top N increased (Fig. 1), suggesting that the overall degree distributions become more consistent across replicates when considering a larger network core.
- **Comparison of Topological Metrics:** While all metrics showed improved stability with larger Top N, Betweenness Centrality often exhibited slightly higher MGP and Fleiss’ Kappa values compared to Degree or Bridging Centrality, suggesting it might be marginally more robust in identifying consistently critical nodes under RIS sampling. Heatmaps visualizing pairwise Jaccard coefficients further illustrated the increasing consistency with larger Top N (Fig. 2).

**Fig. 1.**
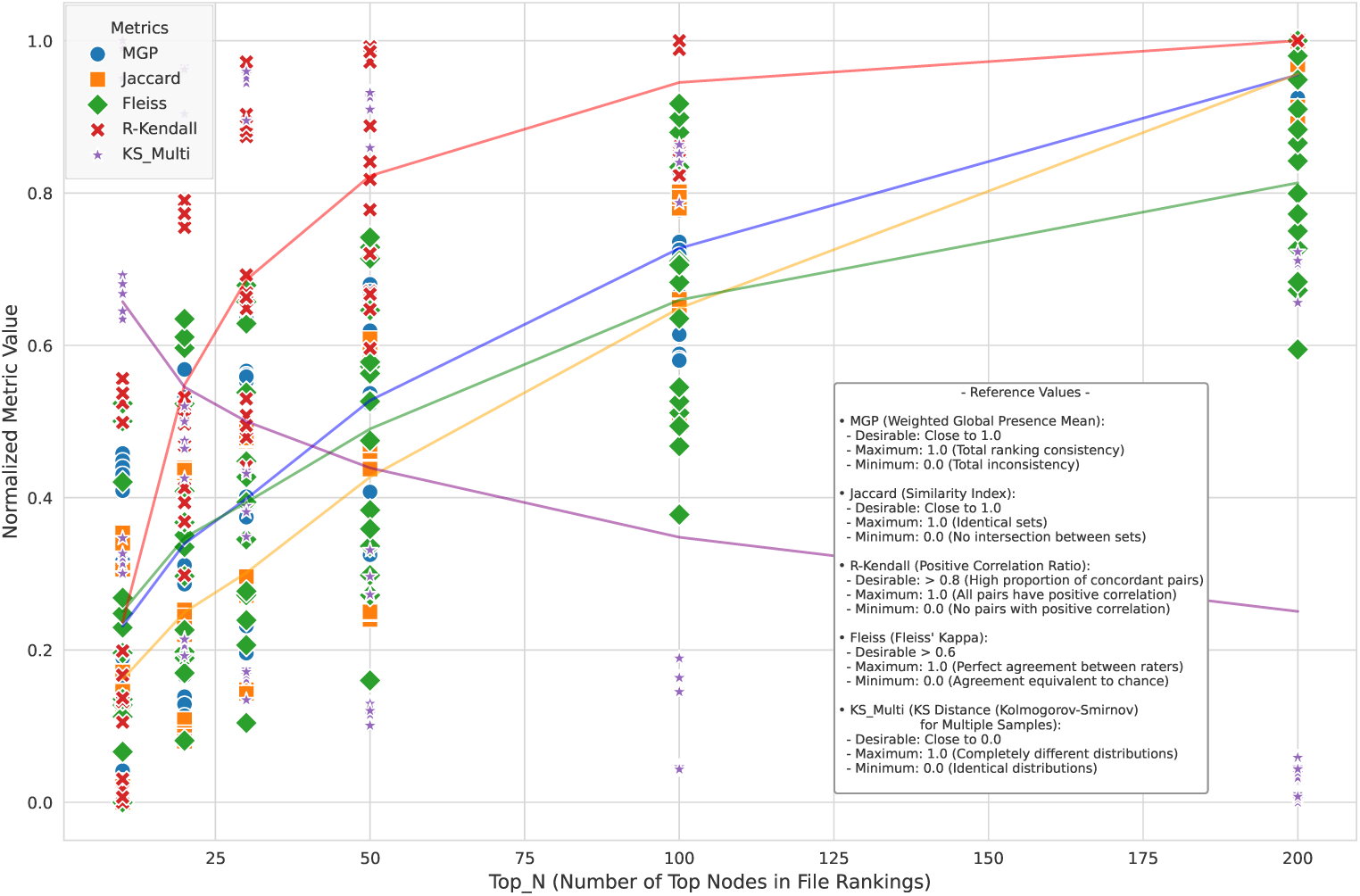
Behavior of Normalized Stability Metrics as a Function of Top N in Machine Learning Mode. Metrics include Weighted Global Presence Mean (MGP), Jaccard Coefficient, Fleiss’ Kappa, Kendall Ratio Positive (R-Kendall), and Multi-Sample KS Distance (KS Multi, inverted scale: lower is better). All metrics are normalized to the range [0, 1] for comparison. Lines connect average values across replicates, showing increased stability (higher MGP, Jaccard, Fleiss, R-Kendall; lower KS Multi) with larger Top N values, indicating RIS preserves core network structure better when analyzing larger node sets.

**Fig. 2.**
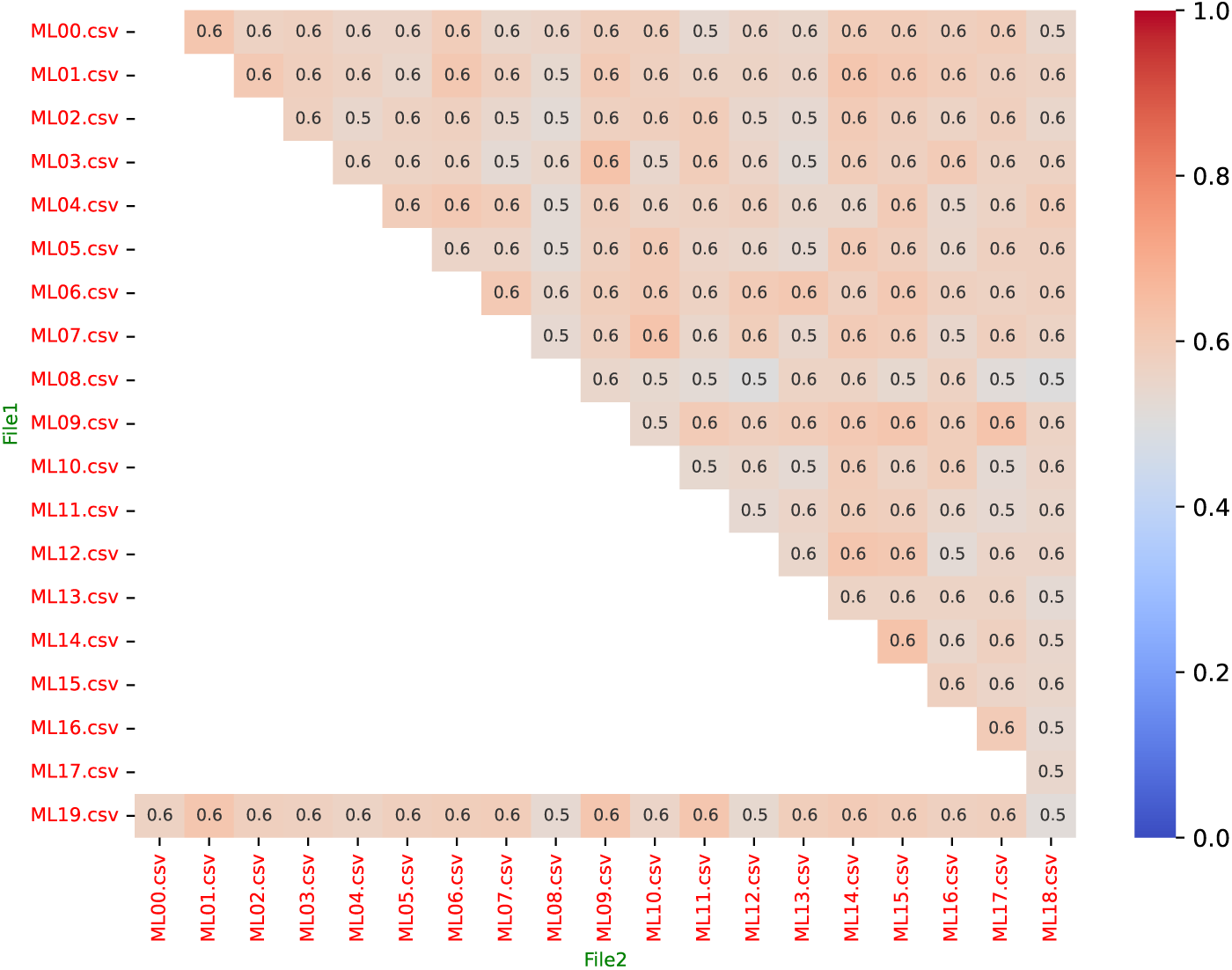
Example Heatmap of Pairwise Jaccard Coefficients for RIS Evaluation. This plot shows the similarity (Jaccard index) between the sets of Top N nodes from different replicate runs (files ML00.csv to ML19.csv from a specific test case: ML mode, Top N=200, 20 files, Degree metric). Axes represent pairs of replicate files. The color scale intensity indicates the Jaccard coefficient value (0.0 to 1.0); in this scheme, blue shades represent low values (near 0.0), and red shades represent high values (near 1.0). Therefore, red shades indicate high consistency between replicates, representing high Jaccard indices (values closer to 1.0). Observing such heatmaps across different Top N values revealed an increased prevalence of these reddish shades (indicating higher consistency) for larger Top N.

**Impact of** FilesNum: Varying the number of input files showed less consistent effects. While Fleiss’ Kappa generally increased with the addition of more files (suggesting a better identification of a common core structure), MGP showed a slight decrease, potentially due to the increased noise or variability introduced by analyzing more diverse datasets simultaneously. Jaccard and Kendall Tau correlations exhibited more erratic behavior relative to the number of files.

In conclusion, the evaluation demonstrates that the RIS algorithm is a viable and effective strategy for managing the computational complexity of PPI network generation in GenPPi, particularly when using the Machine Learning mode. While it introduces variability, especially for small Top N values, the essential topological structure and hierarchical ranking of critical nodes are well-preserved when analyzing a sufficiently large portion of the network core (Top N *≥* 100 recommended for a balance between computational efficiency and structural integrity). The stability in Features mode is near-perfect, validating its use without significant concern for stochastic effects. These findings support the use of RIS within GenPPi 1.5 as a practical approach to enable the analysis of large-scale genomic datasets.

### 3.3 Comparison of Results: GenPPi v1.0 vs. Use of Features and Machine Learning Features

The analysis of the results obtained by GenPPi in version 1.0 compared to incorporating the *Features* and *Machine Learning* functionalities in GenPPi 1.5 reveals a significant improvement in the prediction of protein interaction networks. We observed a consistent trend of increasing the number of nodes and edges, indicating a greater complexity and scope of the networks generated after implementations.

Initially, in version 1.0 of the GenPPi, the genomes of *Buchnera aphidicola* resulted in a relatively low number of nodes and edges, as evidenced in the data in table 3.

**Table 3.** Impact of GenPPi 1.5 Enhancements on Network Size for *Buchnera aphidicola* Genomes. This table compares the number of predicted nodes and edges generated by the original GenPPi (v1.0) with those generated by GenPPi 1.5, utilizing either the ’Features’ mode or the ’Machine Learning’ mode. Results are presented for five different *Buchnera aphidicola* datasets, demonstrating the expansion in network complexity achieved with the new methodologies.

| File | Number of<br>Proteins by<br>Genomes | GenPPi 1 |  | GenPPi 1.5 |  |  |  |
| --- | --- | --- | --- | --- | --- | --- | --- |
|  |  | Nodes | Edges | Features |  | Machine Learning |  |
|  |  |  |  | Nodes | Edges | Nodes | Edges |
| Ba_Bp.dot | 553 | 46 | 91 | 217 | 1746 | 539 | 30220 |
| Ba_Ua.dot | 591 | 111 | 314 | 450 | 6115 | 581 | 40049 |
| Ba_G002.dot | 621 | 120 | 395 | 529 | 9166 | 617 | 41969 |
| Ba_Sg.dot | 620 | 88 | 203 | 430 | 5583 | 614 | 41483 |
| Ba_Ak.dot | 559 | 119 | 339 | 448 | 7917 | 559 | 44005 |
| **Summary** | 2944 | 484 | 1342 | 2074 | 30527 | 2910 | 197726 |

However, after introducing the *Features* and *Machine Learning* features, there was a notable increase in the number of nodes and edges in all the analyzed samples. This expansion of protein interaction networks suggests a greater ability of the improved software to identify and predict relationships between proteins, providing a more comprehensive and detailed view of the underlying biological processes.

The results also indicate an upward trend in the complexity of the generated networks, reflected in a significant increase in the number of nodes and edges. In summary, the comparative results between the different versions of GenPPi demonstrate the benefits of using additional resources in predicting protein interaction networks.

### 3.4 Temporal Efficiency Analysis

To address the computational efficiency of the enhanced GenPPi (v1.5) relative to the original version (v1.0), comparative runtime tests were performed using the *Buchnera aphidicola* genome dataset on a 40-core processor machine, employing a dynamic neighborhood expansion window starting at 3 for all runs. GenPPi v1.0, using its simpler similarity heuristics and serial processing, completed the analysis rapidly in approximately 4 seconds, identifying 156 interactions just for the *Ba ak* genome. GenPPi v1.5 in Features mode, which replaces the simple heuristic with a more complex comparison across 60 biophysical features, took significantly longer (39 seconds) despite utilizing parallel processing. This increase reflects both the higher intrinsic cost of the feature comparison and the larger workload imposed on downstream analyses by the substantially greater number of identified interactions (6,863). The Machine Learning mode of GenPPi v1.5, incorporating the Random Forest classifier and the RIS algorithm, required the longest execution time (117 seconds), yielding the highest number of interactions (45,934). We attributed this further increase to the computational expense of evaluating the 500-tree Random Forest model for similarity prediction and the subsequent processing of the vastly expanded set of potential interactions by the parallelized genomic context algorithms, even with RIS mitigating the combinatorial explosion. These results quantify the expected trade-off: the enhanced sensitivity and broader interaction discovery capabilities of GenPPi 1.5, particularly in ML mode, come at the cost of increased computational time, even when leveraging parallel execution on multi-core hardware.

### 3.5 Comparison with STRING

The data presented below refer to the *Ba ak* genome of the bacterium *Buchenera aphidicola* and were obtained through a SQL query to the *compareba.dump* database. The SQL file *count intersect distinct.sql* contains the definition of a function called *count intersect*. This function returns a table with information about the intersection of different genomic datasets. Function code involves identifying unique elements and their intersections and calculating metrics such as the proportion of unique intersections and interactions between elements. After defining the function, a SELECT statement calls the *count intersect* function and returns the query results.

The conserved phylogenetic profile suggests interactions between pairs of proteins that share identical profiles. Pairing proteins with similar profiles can also lead to interaction. To incorporate this notion of similarity rather than exact identity, the *-ppdifftolerated* parameter in GenPPi establishes the tolerated difference between profiles. The flexibility provided by the *-ppdifftolerated* parameter allows for a more comprehensive analysis of protein-protein interactions, considering the similarity of profiles [5].

Table 4 presents the results of different implementations of GenPPi, varying parameters such as the number of proteins in a phylogenetic profile and the tolerated difference between the profiles of a pair of proteins. Each row in Table 4 represents a specific implementation. The table features various metrics, including *query unique, string intersect unique, unique intersect ratio, and string proportion* for different parameters. For example, the term *tol* is synonymous with the *ppdifftolerated parameter*. The N number accompanying the *RIS* represents the attempts to determine whether the *RIS* algorithm would be triggered when a profile set had more than N proteins. The standard that has been established (*mlris*) is N = 100.

**Table 4.** Comparative Analysis of GenPPi 1.5 Prediction Overlap with STRING Database for the *B. aphidicola* Ba ak Genome. This table evaluates various GenPPi 1.5 implementation configurations (Features mode, Machine Learning mode with RIS, and variations in -ppdifftolerated and RIS N parameters) against the STRING database. Metrics include the number of unique predicted pairs, the overlap with STRING interactions, the ratio of unique predictions to STRING overlap, and the proportion of STRING interactions recovered by GenPPi.

| Implementation | Unique Pairs | STRING Overlap | Unique / Overlap Ratio | STRING Proportion |
| --- | --- | --- | --- | --- |
| *string* | 27041 | 54082<sup>1</sup> | 0.50<sup>2</sup> | 2.00<sup>2</sup> |
| *Features* | 7917 | 2773 | 2.86 | 0.10 |
| *mlris*<sup>3</sup> | **49077** | **16681** | **2.94** | **0.62** |
| *mlristol1*<sup>34</sup> | 103743 | 26924 | 3.85 | 1.00 |
| *mlristol2*<sup>34</sup> | 128579 | 29846 | 4.31 | 1.10 |
| *mlristol3*<sup>34</sup> | 144468 | 32278 | 4.48 | 1.19 |
| *mlris35tol3*<sup>345</sup> | 135617 | 23840 | 5.69 | 0.88 |
| *mlris35tol2*<sup>345</sup> | 114998 | 20034 | 5.74 | 0.74 |

<sup>1</sup>STRING overlap for itself counts directed pairs (A-B, B-A).
<sup>2</sup>Ratio and Proportion for STRING adjusted assuming 27,041 unique interactions.
<sup>3</sup>'mlris' indicates Machine Learning mode with RIS algorithm (default N=100).
<sup>4</sup>'tolN' indicates using the '-ppdifftolerated' parameter N (1, 2, or 3).
<sup>5</sup>'mlrisN' indicates the RIS threshold set to N (here, N=35 is shown).
**Column Definitions:** **Unique Pairs:** Number of unique interaction pairs predicted by GenPPi (A-B/B-A counted once). **STRING Overlap:** Number of unique GenPPi pairs also present in STRING (unique interactions = 27,041). **Unique / Overlap Ratio:** Ratio of Unique Pairs to STRING Overlap; values > 1 indicate predictions beyond STRING. **STRING Proportion:** Recall against STRING (STRING Overlap / 27,041).

In table 4, we have results for the *Features* and the *Machine Learning* modes. Only the sixty traits described in section 2.1 were used with a cut-off factor of twenty-five identical features between a pair of proteins, considering them identical for eventual interaction conservation analyses. The term *ml* is synonymous with the *Machine Learning* mode of GenPPi 1.5. The *ml* mode is triggered when the *Features* mode concludes that, based on the lack of at least twenty-five identical features among a pair of proteins, a pair of proteins is not identical.

The *query unique* column represents the number of implementation-related individual interactions. In comparison with STRING, column *string intersect unique* indicates the number of unique interactions that match those of STRING, while column *unique intersect ratio* represents the ratio of the number of unique interactions in GenPPi to STRING. The *string proportion* column indicates the proportion of interactions in GenPPi that are also present in STRING.

Note duplicated values referring to the row corresponding to the STRING because the data source in the STRING considers the interaction between A and B different from B and A. There is no duplication in the other data. Therefore, the actual value of *unique intersect ratio* and *string proportion* in the line corresponding to the STRING is 1 and 1, instead of 0.5 and 2, respectively. The reason is that the *count intersect* function is intended to obtain metrics for implementations made in GenPPi and not for STRING.

*Unique intersect ratio* is a crucial metric for understanding the effectiveness of different GenPPi implementations. This ratio represents the number of predicted interactions divided by the number of interactions intersecting with the STRING.

Therefore, for every interaction that matches the STRING, many other interactions do not belong to the STRING. We can interpret the existence of interactions absent in STRING as an inverted distance measure of the STRING, where a *unique intersect ratio* of 1 would indicate a perfect match with the STRING. The higher the *unique intersect ratio* value, the greater the difference between the phylogenetic profiles of a pair of proteins is, allowing them to be classified as interacting, which may result in more predicted interactions by GenPPi. However, some of these additional interactions may not be present in STRING. These interactions, absent from STRING but supported by GenPPi’s analysis of conserved genomic features, represent putative candidates for discovery. Interactions predicted by GenPPi in STRING can be useful for discovering potential new interactions, but can also lead to more false positives. Therefore, it is important to strike a balance when adjusting the *unique intersect ratio* value.

It is important to consider that a high average degree of edges, determined by the value of *unique intersect ratio*, can hinder a topological analysis, as it generates many interactions. Therefore, when evaluating the effectiveness of different GenPPi implementations, it is crucial to consider not only the STRING match but also the structure and topology of the resulting network.

As the value of *tol* increases (from *tol1* to *tol3* ), the *unique intersect ratio* also increases. Greater values to *tol* suggest that allowing a greater difference between phylogenetic profiles may result in greater agreement with the STRING. Similarly, increasing the number of proteins in a profile (from *mlris10* to *mlris35* ) also appears to increase *unique intersect ratio*. However, it is important to note that *unique intersect ratio* higher does not necessarily indicate a better prediction of protein-protein interactions. While greater agreement with the STRING may suggest greater accuracy, it is also possible that some valid interactions may be lost. Therefore, it is crucial to consider the balance between sensitivity (the ability to identify interactions correctly) and specificity (the ability to exclude non-interactions correctly) when evaluating the effectiveness of different GenPPi implementations.

In addition, the *string proportion* represents the result of dividing the number of single-intersect interactions of the specific GenPPi implementation by the number of STRING single-intersecting interactions (i.e., 27041), a measure of the proportion of interactions in GenPPi that are also present in STRING. For the implementation using *Machine Learning* and the *RIS* algorithm *mlris* it achieved a percentage of 62%. For *mlris35tol3* and *mlris35tol2*, a percentage of 88% and 74% respectively. For *mlristol1*, *mlristol2* and *mlristol3*, values higher than 100%, being respectively 100%, 110% and 119%. Values higher than 100% occur in certain cases where the adopted implementation recognizes the STRING values of interactions repeated in different scenarios.

Among the various implementations, the most significant is *mlris*, which utilizes machine learning and RIS algorithms, generating a total of 49,077 unique interactions, representing a proportion 1.81 times greater than the 27,041 unique interactions present in STRING. Of this total, 16,681 interactions coincide with STRING, representing 62% of the interactions predicted in STRING. The ratio between the number of unique interactions of the implementation divided by the number of interactions that intersect with the STRING was 2.94, indicating that for each coincident interaction, there is 1.94 interaction that does not match the STRING. These quantitative results demonstrate the ability of the GenPPi program, equipped with the RIS algorithm, to generate predictions of protein interactions with a significant proportion of overlap with the interactions documented in STRING while presenting a network that is not extremely dense.

The presented results demonstrate significant improvements in GenPPi’s capabilities. However, they also highlight important trade-offs and require careful interpretation within the broader context of PPI prediction.

## 4 Discussion

### 4.1 Interpreting Predictions and Network Density

The protein interactions predicted by GenPPi that are not present in the STRING database do not necessarily imply the non-existence of these interactions or the incorrectness of the GenPPi software. The correctly implemented algorithms for conserved phylogenetic profile and gene neighborhood traits conferred between the GenPPi versions speak in favor of a high probability that these interactions occur. However, an overly dense protein network can make it difficult to identify significant differences in the network topology, hindering deeper analyses. On the other hand, an overly sparse network is also not ideal. Therefore, the search for an intermediate-density network becomes a relevant objective.

### 4.2 Performance Comparison: Features vs. Machine Learning Modes

Comparing the operational modes reveals significant differences in Table 4. The Features mode, operating without machine learning, generates a considerably smaller number of unique interactions (‘query unique‘). In contrast, the Machine Learning mode produces substantially larger networks. This difference impacts the comparison with STRING: while ML implementations achieve higher overlap (‘string proportion‘ reaching up to 1.19, indicating the capture of nearly all STRING interactions and potentially identifying duplicates present in STRING itself), the Features mode shows lower overlap (e.g., 0.10).

However, the increased capture rate in ML mode brings together a much larger number of predicted interactions not found in STRING, as shown by the higher ‘unique intersect ratio‘ values (reaching up to 5.74). Consequently, the ML-generated networks exhibit significantly higher edge density due to many unique predictions (‘query unique‘), which can complicate subsequent topological analyses compared to the sparser networks generated from the Features mode. A clear trade-off between sparse and dense nets: the sparser Features network may be more amenable to topological inference while potentially missing some known interactions (lower ‘string proportion‘). Conversely, the denser ML network captures more known biology (higher ‘string proportion‘) but poses greater challenges for analysis due to its complexity and the higher ratio of novel-to-known predictions (higher ‘unique intersect ratio‘).

### 4.3 Validation Against STRING Database

It is crucial that the predicted protein networks significantly overlap with the protein interactions documented in the STRING database. String overlapping validates the accuracy of the forecasts and strengthens the reliability of the results. The results suggest that the interactions generated by GenPPi significantly overlap with those predicted by STRING, which is considered a gold standard. The interpretation of these data suggests that GenPPi can capture substantial interactions similar to those of STRING, thereby validating the software’s effectiveness and relevance in generating protein-protein interaction networks. This quantitative comparison offers valuable insights into the quality and accuracy of the interactions generated by GenPPi in comparison to the standard set by STRING.

### 4.4 Utilizing Interaction Confidence Scores

An interaction can also have a high probability of being true or indicate a lower probability of being a real interaction. To work around the problem of the high number of interactions, some parameters can be adjusted to increase the confidence of the probability of an interaction. GenPPi and STRING provide information on known interactions and predictions of protein interactions. STRING, for example, assigns a confidence value or a confidence score to indicate the strength or probability of the interaction. This value is calculated based on various data sources, including experimental evidence, gene coexpression, and sequence similarity. We displayed the confidence value assigned by STRING and GenPPi as a number between 0 and 1.

### 4.5 Overall Performance and Non-Randomness of Predictions

The results obtained through using GenPPi demonstrate its usefulness in improving the quality and efficiency of investigations. A key step contributing to this achievement was the RF model demonstrating high average specificity ( 90%) but moderate average sensitivity (57.3%), resulting in an average F1-Score of 51%. Such a moderate F1-score highlights the model’s prioritization of minimizing false positives. Another important point to discuss is the number of edges presented by the GenPPi interaction networks. The average number of proteins in our set of five *Buchnera aphidicola* genomes is about 590. All possible combinations of protein pairs, valid or not as interacting, are about 348,100 per genome. Considering our best outcome in Table 4, genome *mlris* and *string proportion* 0.62, among all possibilities of interactions, we gathered 92,137 or just 26% of the odds, considering the *Machine Learning* mode. When we configure GenPPi with *Features* mode, the slice possible protein pairs from the odds is only 2%. Even in the worst GenPPi configuration scenario, genome *mlristol3*, we obtained 0.41% of the odds. However, we obtained 100% of the STRING predictions, followed by the configuration genome *mlristol1*, getting a lower slice from the odds of 29% of unique interactions. These numbers evidence that the GenPPi interaction networks are not pure by chance. Despite some configurations yielding a higher number of candidate interactions, we rely on a significant number of interactions predicted by other software and for trustworthy algorithms.

### 4.6 GenPPi’s Niche and Distinct Advantages

While GenPPi’s predictive methodology, based on genomic context, shares principles with comprehensive databases like STRING (a significant portion of whose predictions are also computational), it offers several distinct advantages, particularly for discovery-driven research and analysis of less-characterized organisms. Firstly, GenPPi facilitates the de novo prediction of protein-protein interaction (PPI) networks for any set of user-provided bacterial or archaeal genomes. This is crucial for newly sequenced organisms or specific strains not yet incorporated into public databases, where methods relying on pre-existing annotations or interolog mapping for novel proteins may fail if suitable homologs with known interactions are absent. Secondly, the software enables customizable, large-scale comparative analyses across multiple genomes locally, offering flexibility beyond typical web-server constraints and allowing for investigations tailored to specific evolutionary or functional questions. Thirdly, GenPPi operates independently of 3D structural information, providing a valuable complementary approach to structure-based prediction methods, which are often limited by the availability of high-quality structural data on a proteome-wide scale [3, 6].

While direct outperformance of established, curated databases like STRING across all metrics is not the primary claim for this version, GenPPi’s strength lies in its accessibility for novel genome analysis, its capacity for custom network generation, and its foundational ab initio approach that can propose interactions even for unannotated proteins, thereby serving as a powerful hypothesis generation tool.

### 4.7 Rationale for Random Forest Selection

While a wide array of machine learning algorithms, such as XGBoost or KNN, could potentially be applied to identify protein homology, a comprehensive benchmark against alternatives was not the primary focus of this work, nor would it necessarily reflect the specific constraints and optimization goals inherent to the GenPPi pipeline. The primary objective was to enhance GenPPi’s ability to efficiently predict proteinprotein interactions (PPIs) within its existing architecture, which heavily relies on the subsequent, computationally intensive analysis of conserved phylogenetic profiles and gene neighborhoods.

Although increasing the recall of potential protein similarities is desirable, naively maximizing the number of predicted similarities exacerbates the computational burden of these downstream steps. GenPPi’s core innovation lies in identifying similarities and intelligently filtering and prioritizing them based on genomic evidence to generate a manageable and biologically meaningful network. Simply using a classifier with the highest possible recall could overwhelm this filtering process.

The implementation of the RIS algorithm directly addresses this challenge. It allows leveraging the sensitivity of Random Forest to identify potential interactions while mitigating the prohibitive computational cost associated with analyzing an excessive number of candidates. Preliminary experiments with alternative algorithms indicated that some achieved slightly higher initial recall. However, considering that our main goal was not to create a machine learning benchmark, we opted not to use those, thereby avoiding the need to modify our source code dramatically.

Furthermore, Random Forest offered a practical advantage due to the availability of the robust cl-random-forest library implemented in Common Lisp, the same language used for GenPPi’s development. A native Common Lisp software facilitated seamless integration and leveraged the strengths of the Lisp environment. GenPPi, combining Random Forest’s balanced performance with the RIS reduction strategy, achieves prediction rates characterized by moderate specificity ( 90%) and moderate sensitivity ( 57%), which are sufficient to generate high-quality PPI networks within the practical constraints of its analytical pipeline, striking a necessary balance between sensitivity and computational feasibility.

The decision to employ Random Forest (RF) was also informed by several practical and strategic considerations within the GenPPi development context. While Deep Neural Networks (DNNs), including Convolutional Neural Networks (CNNs) and Graph Neural Networks (GNNs), have demonstrated remarkable performance in various bioinformatics tasks, including PPI prediction [6–9], their integration posed challenges that RF helped to mitigate for this version of GenPPi. A key factor was the development environment and ease of integration. GenPPi is implemented in Common Lisp, and the availability of the cl-random-forest library [19] a high-performance, native

Common Lisp implementation allows for seamless and efficient integration without introducing complex dependencies on external Python-based DNN frameworks or extensive inter-language communication overhead. This streamlined the development and deployment process. Furthermore, our experience in a related domain-predicting non-classical protein secretion using biophysical features similar to those in GenPPi showed that a well-optimized RF model could achieve predictive accuracy comparable to more complex machine learning architectures but with significantly faster classification times and a more straightforward training process [29]. In Oliveira et al. (2024), an RF classifier achieved approximately 91% accuracy in classifying non-classical secreted proteins, demonstrating RF’s capability with amino acid propensity-derived features. Given that GenPPi’s RF model acts as an initial filter for a vast number of potential protein pairs, this computational efficiency is advantageous. While a direct, exhaustive benchmark against various DNN architectures for the specific task within GenPPi was beyond the scope of this work, the chosen RF approach provides a robust and scalable solution. We acknowledge the strong potential of DNNs, and exploring their application within GenPPi remains a promising avenue for future development. However, for the current GenPPi 1.5, RF offered an optimal trade-off between predictive performance, computational tractability within our pipeline, ease of integration, and development agility.

### 4.8 Validation of Novel Interactions

Beyond the choice of classifier, a critical aspect concerns the validation of interactions predicted by GenPPi, particularly those not found in reference databases like STRING. A crucial aspect of PPI prediction involves identifying novel interactions that have not been previously documented. However, the experimental validation of computationally predicted interactions presents significant challenges common to all large-scale prediction methods. It is worth noting that reference databases, such as STRING, while invaluable, contain many predicted interactions that lack direct experimental validation. The number of novel interactions proposed by methods like GenPPi (or, indeed, by established databases like STRING) can be considerable. No highthroughput computational method exists to empirically validate them biologically *en masse*.

GenPPi and databases like STRING rely heavily on computational predictions, with only a fraction of interactions typically having direct experimental support. Therefore, protein interactions predicted solely by GenPPi, absent in STRING, should not be automatically dismissed as false positives. These represent putative interactions, supported by rigorous genomic evidence analyzed by GenPPi and rules validated across multiple GenPPi versions. They serve as prioritized candidates for future experimental investigation, potentially leading to new biological insights, particularly in less-studied organisms where databases like STRING may have limited coverage.

The goal of GenPPi is not necessarily to perfectly replicate existing databases but to leverage genomic context to propose biologically plausible networks, including novel interactions that warrant further study. While striving for significant overlap with known interactions, as demonstrated by the 62% overlap with STRING, validates the overall approach. The novel predictions highlight the discovery potential inherent in *ab initio* genome analysis. Focusing solely on replicating known data would limit the tool’s utility for exploring the unknown biology of diverse organisms. Comprehensive experimental validation remains a resource-intensive bottleneck for the entire field, lying beyond the scope of the computational prediction tool itself.

### 4.9 Open Source Availability

Using open-source code in all versions of GenPPi presents numerous advantages for scientific research. Access to open-source code provides benefits such as customization, control, cost reduction, collaboration, transparency, and reliability. However, it is essential to note that scientific sources, such as STRING, are not publicly accessible, which limits access to the source code and compromises the transparency and reliability of the results. GenPPi version 1.5 is available for download at the following address: https://genppi.facom.ufu.br/

### 4.10 User’s rules

A key aspect of the GenPPI software is its ability to transfer the decision about how many and which genomes to use in constructing the protein interaction network to the end user. All the code was written in Common Lisp and compiled using SBCL, allowing the creation of complex protein networks predicted from tens or hundreds of genomes. GenPPi stands out for its versatility and design, which makes it compatible with various operating systems and hardware configurations. We optimized program versions to meet different needs and usage scenarios. It is easy to access and use, requiring no specific expertise in computing or biology. We designed the software to facilitate the discovery of novel relationships between proteins and their associated biological processes.

## 5 Conclusion

Based on the results obtained with GenPPi, it is possible to infer that the software offers functionalities that are valuable for molecular biology and bioinformatics researchers. GenPPi’s ability to generate protein interaction networks from specific, unpublished genomic data without relying on external databases and with validated predictive performance enables researchers to explore protein interactions in lesser-studied genomes or organisms with unique characteristics, thereby broadening the scope of scientific investigations. While direct experimental validation of all novel predictions remains a field-wide challenge, GenPPi provides candidates prioritized by strong genomic evidence. The results of the latest version of GenPPi demonstrate a significant advance in protein interaction studies.

## 6 Availability of software

- **Project name:** GenPPi
- **Project home page:** https://genppi.facom.ufu.br/, https://github.com/ santosardr/genppi
- **Operating system(s):** Platform independent for source code. Pre-compiled binaries are provided for Linux, Microsoft Windows, and macOS.
- **Programming language:** Common Lisp (SBCL).
- **Other requirements:** Steel Bank Common Lisp (SBCL) for compiling from source. The ‘cl-random-forest‘ Quicklisp library is required to train custom RF models.
- **License:** MIT License. The software is freely available as open source.
- **Any restrictions to use by non-academics:** None. The software is freely available for both academic and non-academic use under the terms of the MIT License.

The source code for GenPPi version 1.5 is available on GitHub: https://github.com/santosardr/genppi. This repository includes the source code and pre-compiled binaries. The Random Forest model training scripts are available at https://github.com/santosardr/non-CSPs/tree/main/src/lisp. Scripts and instructions for users to train custom models (‘src/tools/makearff.bash‘ and ‘src/tools/crafting-RF-training-model.lisp‘) are also provided in the GenPPi GitHub repository.

## 7 Declarations

### Ethics approval and consent to participate

Not applicable.

### Consent for publication

Not applicable.

### Availability of data and material

The datasets generated and/or analyzed during the current study are available within the article and its cited repositories. Specifically, the ‘compareba.dump‘ file for the *Buchnera aphidicola* analysis is available in the GenPPi GitHub repository at https://github.com/santosardr/genppi. Detailed RIS evaluation data are available in the Zenodo repository, https://doi.org/10.5281/zenodo.10452237 [28]. Training and test sets for the Random Forest model are available from the corresponding author upon reasonable request. The software itself is available as detailed in the ”Availability of software” section.

### Authors’ contributions

ASi, CM, IG, LSi, MP, MB, and NA contributed to various aspects of the software development, data generation, and data analysis. ARS conceived the project, supervised the work, developed core components of the software, and wrote the manuscript. All authors read and approved the final manuscript.

## Acknowledgements

We thank the Federal University of Uberl^andia (UFU) for providing technical support and web server facilities.

## Clinical trial registration

Not applicable.

## References

[1] Tang, T., Zhang, X., Liu, Y., Peng, H., Zheng, B., Yin, Y., Zeng, X.: Machine learning on protein-protein interaction prediction: models, challenges and trends. Briefings in Bioinformatics 24, 1–11 (2023) 10.1093/BIB/BBAD076

2. Hu, L., Wang, X., Huang, Y.A., Hu, P., You, Z.H.: A Novel Network-Based Algorithm for Predicting Protein-Protein Interactions Using Gene Ontology. Frontiers in Microbiology 12 (2021) 10.3389/FMICB.2021.735329/BIBTEX

[3] Sunny, S., Jayaraj, P.: Protein–protein docking: Past, present, and future. The protein journal 41(1), 1–26 (2022)

[4] Snider, J., Kotlyar, M., Saraon, P., Yao, Z., Jurisica, I., Stagljar, I.: Fundamentals of protein interaction network mapping. Molecular Systems Biology 11(12), 848 (2015) 10.15252/msb.20156351

[5] Anjos, W.F., Lanes, G.C., Azevedo, V.A., Santos, A.R.: GENPPI: standalone software for creating protein interaction networks from genomes. BMC Bioinformatics 22, 1–26 (2021) 10.1186/S12859-021-04501-0/TABLES/5

[6] Song, B., Luo, X., Luo, X., Liu, Y., Niu, Z., Zeng, X.: Learning spatial structures of proteins improves protein–protein interaction prediction. Briefings in Bioinformatics 23(2) (2022) 10.1093/bib/bbab558

7. Soleymani, F., Paquet, E., Viktor, H., Michalowski, W., Spinello, D.: Proteinprotein interaction prediction with deep learning: A comprehensive review. Computational and Structural Biotechnology Journal 20, 5316 (2022) 10.1016/J.CSBJ.2022.08.070

[8] Gao, Z., Jiang, C., Zhang, J., Jiang, X., Li, L., Zhao, P., Yang, H., Huang, Y., Li, J.: Hierarchical graph learning for protein–protein interaction. Nature Communications 14(1), 1234 (2023) 10.1038/s41467-023-36736-1

[9] Tang, T., Li, T., Li, W., Cao, X., Liu, Y., Zeng, X.: Anti-symmetric framework for balanced learning of protein–protein interactions. Bioinformatics 40(10) (2024) 10.1093/bioinformatics/btae603

[10] Zhang, W., Xue, X., Xie, C., Li, Y., Liu, J., Chen, H., Li, G.: CEGSO: Boosting Essential Proteins Prediction by Integrating Protein Complex, Gene Expression, Gene Ontology, Subcellular Localization and Orthology Information. Interdisciplinary sciences, computational life sciences 13, 349–361 (2021) 10.1007/S12539-021-00426-7

11. Oliveira, G.S., Santos, A.R.: Using the Gene Ontology tool to produce de novo protein-protein interaction networks with IS A relationship. Genetics and molecular research : GMR 15 (2016) 10.4238/GMR15049273

[12] Szklarczyk, D., Franceschini, A., Wyder, S., Forslund, K., Heller, D., HuertaCepas, J., Simonovic, M., Roth, A., Santos, A., Tsafou, K.P., Kuhn, M., Bork, P., Jensen, L.J., Mering, C.: STRING v10: protein-protein interaction networks, integrated over the tree of life. Nucleic acids research 43, 447–452 (2015) 10.1093/nar/gku1003

13. Kang, Y., Wang, X., Xie, C., Zhang, H., Xie, W.: BBLN: A bilateral-branch learning network for unknown protein-protein interaction prediction. Computers in Biology and Medicine 167, 107588 (2023) 10.1016/J.COMPBIOMED.2023.107588

[14] Sardiu, M.E., Gilmore, J.M., Groppe, B.D., Dutta, A., Florens, L., Washburn, M.P.: Topological scoring of protein interaction networks. Nature Communications 2019 10:1 10, 1–14 (2019) 10.1038/s41467-019-09123-y

[15] Kawashima, S., Pokarowski, P., Pokarowska, M., Kanehisa, M.: AAindex: amino acid index database. Nucleic acids research 28(1), 374–374 (2000) https://doi. org/10.1093/nar/28.1.374

[16] Jia, J., Liu, Z., Xiao, X., Liu, B., Chou, K.-C.: Identification of protein-protein binding sites by incorporating the physicochemical properties and stationary wavelet transforms into pseudo amino acid composition. Journal of Biomolecular Structure and Dynamics 34(9), 1946–1961 (2016)

[17] Breiman, L.: Random forests. Machine learning 45(1), 5–32 (2001) 10.1023/A:1010933404324

[18] Speiser, J.L., Miller, M.E., Tooze, J., Ip, E.: A comparison of random forest variable selection methods for classification prediction modeling. Expert Systems with Applications 134, 93–101 (2019) 10.1016/J.ESWA.2019.05.028

19. Imai, S.: CL-random-forest: Random forest in Common Lisp. GitHub (2024). https://github.com/masatoi/cl-random-forest

[20] Ren, S., Cao, X., Wei, Y., Sun, J.: Global refinement of random forest. Proceedings of the IEEE Computer Society Conference on Computer Vision and Pattern Recognition 07-12-June-2015, 723–730 (2015) 10.1109/CVPR.2015.7298672

21. Bastian, M., Heymann, S.: Gephi : An Open Source Software for Exploring and Manipulating Networks. ICWSM 8, 361–362 (2009) 10.1136/ qshc.2004.010033

[22] Camacho, C., Coulouris, G., Avagyan, V., Ma, N., Papadopoulos, J., Bealer, K., Madden, T.L.: BLAST+: architecture and applications. BMC bioinformatics **10**(1), 1–9 (2009) 10.1186/1471-2105-10-421

[23] Pereira, G., Ghosh, P., Santos, A.: A Bridging Centrality plugin for GEPHI and a case study for Mycobacterium tuberculosis H37Rv. IEEE/ACM Transactions on Computational Biology and Bioinformatics 18(6), 2741–2746 (2021)

[24] Jaccard, P.: Etude comparative de la distribution florale dans une portion des Alpes et des Jura. Bull Soc Vaudoise Sci Nat 37, 547–579 (1901)

[25] Fleiss, J.L.: Measuring nominal scale agreement among many raters. Psychological bulletin 76(5), 378 (1971)

[26] Kendall, M.G.: A new measure of rank correlation. Biometrika 30(1-2), 81–93 (1938)

[27] An, K.: Sulla determinazione empirica di una legge didistribuzione. Giorn Dell’inst Ital Degli Att 4, 89–91 (1933)

[28] Silva, A., Marquez, C., Godoy, I., Silva, L., Prado, M., Beppler, M., Avila, N., Santos, A.: GenPPI RIS Algorithm Evaluation Dataset. Zenodo. Version 1.5 (2024). 10.5281/zenodo.15098293.

[29] Oliveira, L., Lanes, G., Santos, A.: Enhancing omics analyses of bacterial protein secretion via non-classical pathways. Neural Computing and Applications 36(27), 17045–17055 (2024) 10.1007/s00521-024-09993-4

